# Assessing complex member composition and insecticide resistance status of *Anopheles gambiae s.l.* in three coastal health districts of Côte d’Ivoire

**DOI:** 10.1101/2024.01.10.575138

**Authors:** Jackson IK Kouamé, Constant VA Edi, Julien BZ Zahouli, Ruth MA Kouamé, Yves AK Kacou, Firmain N Yokoly, Constant GN Gbalegba, David Malone, Benjamin G Koudou

**Affiliations:** Unité de Formation et de Recherche Sciences de la Nature, Université Nangui Abrogoua, Abidjan, Côte d’Ivoire; Environnement et Santé, Centre Suisse de Recherches Scientifiques en Côte d’Ivoire, Abidjan, Côte d’Ivoire; Centre d’Entomologie Médicale et Vétérinaire, Université Alassane Ouattara, Bouaké, Côte d’Ivoire; École Supérieure d’Agronomie, Institut National Polytechnique Félix Houphouët Boigny, Yamoussoukro, Côte d’Ivoire; Unité de lutte antivectorielle, Programme National de Lutte Contre le Paludisme, Abidjan, Côte d’Ivoire; Innovative Vector Control Consortium, Liverpool School of Tropical Medicine, Liverpool, United Kingdom

**Author notes:** Corresponding author /.

**Keywords:** *Anopheles gambiae s.l.*, Insecticides resistance, Chlorfenapyr, clothianidin, Coastal Côte d’Ivoire

## Abstract

Although malaria is endemic in Coastal Côte d’Ivoire, updated data on the resistance profile of the main vector, *Anopheles gambiae sensu lato* (*s.l*.), are still lacking, thus compromising decision-making for an effective vector control intervention. This study investigated the complex members and the insecticide resistance in the *Anopheles gambiae s.l.* populations in coastal Côte d’Ivoire. Between 2018 and 2020, bioassays were conducted on female *An. gambiae s.l.* mosquitoes in three coastal health districts (Aboisso, Jacqueville and San Pedro) of Côte d’Ivoire. Pyrethroids deltamethrin, permethrin and alphacypermethrin (1X, 5X and 10X), clothianidin and synergist piperonyl butoxide (PBO) combined with pyrethroid 1X were tested using WHO tube bioassays. Chlorfenapyr was evaluated using CDC bottle bioassays. *An. gambiae* complex members and *Vgsc* 995F, *Vgsc* 995S and Ace-1 280S mutations were identified using polymerase chain reaction (PCR) technique. Overall, *Anopheles. gambiae s.l.* populations were primarily composed of *Anopheles coluzzii* (88.24%, n = 312), followed by *Anopheles gambiae sensu stricto* (*s.s.*) (7.56%) and hybrids (4.17%). These populations displayed strong resistance to pyrethroids at standard diagnostic doses, with mortality remaining below 98% even at 10X doses, except for alphacypermethrin in Aboisso. Pre-exposure to PBO significantly increased mortality but did not fully restore susceptibility, except for alphacypermethrin in Jacqueville. Clothianidin restored full susceptibility in Jacqueville and San Pedro, while chlorfenapyr restored susceptibility in Aboisso at 100 µg ai/bottle and all three districts at 200 μg ai/bottle. *Vgsc* 995F mutation dominated, with frequencies varying from 71.2% to 79.3%. *Vgsc* 995S had low, and showed frequencies ranging from 2.3% to 5.7%. Ace-1 280S prevalence varied between 4.2% and 42.9%. Coastal Côte d’Ivoire’s *An. gambiae s.l.* populations were mainly composed of *Anopheles coluzzii* and showed high resistance to pyrethroids. Clothianidin, chlorfenapyr, and PBO with pyrethroids increased mortality, indicating their potential use as an alternative for malaria vector control.

## Introduction

The World Health Organization (WHO) has reported recently that there were 249 million cases and 608,000 deaths from malaria in 2022 [1]. Malaria is most prevalent in sub-Saharan Africa, where the majority of cases and deaths occur. Within sub-Saharan Africa, countries such as Nigeria, the Democratic Republic of the Congo, and Mozambique have some of the highest malaria burdens. Malaria is one of the deadliest public health diseases, particularly in sub-Saharan Africa among children less than 5 years old. The WHO African region accounted for 95% of global malaria cases and 96% of deaths [2]. More than US$ 3.5 billion have been invested in 2021 for malaria, with a third of this investment (around US$ 1.1 billion) spent by the governments of malaria-endemic countries [2]. The WHO recommended malaria prevention strategies, including the effective vector control tool that have major impacts in reducing the global burden of this disease. The WHO Global Technical Strategy (GTS) aims to reduce malaria incidence and mortality by at least 75% by 2025 and 90% by 2030 which seems to be challenging [2]. In 2021, 242 million Artemisinin-based Combination Therapies (ACTs) were distributed to the public health sector by National Malaria Control Programs (NMCPs) and about 590 million Insecticide Treated Nets (ITNs) were delivered to communities in the period 2019–2021 [2]. To achieve these goals, GTS has called for the development of new vector control tools that must incorporate new insecticide molecules (e.g., clothianidin and chlorfenapyr), synergists (e.g., pyperonyl butoxide: PBO) and/or insecticide mixtures containing at least two active ingredients with different modes of action to mitigate malaria vector resistance to insecticides [2–4]. Côte d’Ivoire’s population lives in high malaria risk areas [5]. In 2021, malaria burden cases in the country were estimated at 7,443,146 cases and 14,906 deaths [2]. The main objectives of the National Control Malaria Program (NMCP) are to reduce malaria morbidity by 40 percent and by 33 percent the malaria mortality in high-burden by 2026 compared to the 2015 baseline [6]. The vector control strategy implemented by the NMCP is mainly based on the mass deployment of long-lasting insecticidal nets (LLINs) every three years and recently by implementing indoor residual sprays (IRS) [7–9] in one district. In Côte d’Ivoire, the National Malaria Strategic Plan 2016–2020 has prioritized indoor residual spraying (IRS) as an additional vector control method to reduce malaria morbidity and mortality [10]. The transmission of malaria in Côte d’Ivoire are ensured by *An. gambiae s.l., An. funestus s.l.* and *An. nili* [7,11]. Species from the *An. gambiae s.l.* complex (*An. gambiae s.s.*, *An. coluzzii*, and hybrids) have been identified across the country [7,12–15]. However, Ivorian *An. gambiae s.l.*, populations exhibit strong resistance to many traditional classes of insecticides (i.e., DDT, pyrethroid, carbamate, and organophosphate) [7,13–17]. Local *An. gambiae s.l.* resistance to insecticides is a severe challenge to the NMCP efforts because all LLINs distributed in Côte d’Ivoire before 2021 were treated only with pyrethroid [7,10]. Therefore, it is urgent to develop effective alternative tools for the sustainable control of insecticide-resistant malaria vectors. So, since 2018, the National Malaria Strategic Plan (NMSP) supported by the U.S. President’s Malaria Initiative (PMI) project has contributed to the generation of insecticide resistance data to conduct a stratification of vector control interventions across the country. From 2018 to 2022, the NMCP and PMI deployed IRS in the health districts of Nassian and Sakassou using clothianidin-based insecticides that resulted in effective malaria vector control [6,7]. Insecticide susceptibility tests were also carried out in order to have nets incorporated with pyrethroids and other active ingredients (synergists, pyrroles, etc.) and with dual active ingredients, for the next LLIN distribution campaign, in line with the entomological stratification data [6,7]. Recently, pyrethroids in combination with either an insect growth regulator, a pyrrole, or a synergist that inhibits the primary metabolic mechanism of pyrethroid resistance within mosquitoes were used in LLINs [3,4,7]. Therefore, in the current study, insecticide resistance status and complex members of *An. gambiae s.l.* were assessed in three coastal health districts of Côte d’Ivoire, namely Aboisso, Jacqueville and San Pedro. The health district of Jacqueville records one of the highest malaria incidences three hundred sixty-seven per mile (367‰), while a lower incidence is reported in the health district of San Pedro (127.1‰) [18,19]. In Côte d’Ivoire, the coastal areas are characterized by a massive concentration of people [20]. These areas are the heart of the economic development of the country and have two main seaports, one in Abidjan and one in San-Pedro. This coastal region is home to the largest number of agro-industrial units, traditional agriculture and smallholdings [11,14,21]. The industrial and traditional crops (e.g., cocoa, rubber, oil palm, coffee, rice) [11,12,14,15,21] are intensively treated with pesticides and insecticides to protect crops [14]. The local *Anopheles* malaria vectors are constantly under strong pressure from insecticides and chemicals, which can lead to the development of their resistance and increased their survival. Therefore, an urgent need is to develop alternative insecticides and evaluate their effectiveness against malaria vectors in resistant areas, mainly on the coastline. The current study is part of a program designed to explore the landscape of malaria *Anopheles* vectors in coastal areas and inform disease control programs of Côte d’Ivoire.

## Methodology

### Study sites

The study was conducted in three health districts, namely Aboisso (latitude 5° 28’ 04” N and longitude 3° 12’ 25” W), Jacqueville (latitude 5° 12’ 02” N and longitude 4° 24’ 44” W) and San Pedro (latitude 4° 44’ 5” N and longitude 6° 38’ 10” W), located in the southern littoral area of Côte d’Ivoire (Fig 1). The local climate is characterized by two major seasons: the long rainy season from April to July, with the peak of rainfalls in June, while the long dry season extends from October to March. The annual rainfall average is 1848 mm with an annual temperature average of 27 °C. In the three health districts, economic activities were dominated by agriculture leading to large agricultural areas for food crops and cash crops (rubber, oil palm, pineapple, cassava, yam, banana) [20,22,23]. Farmers use several types of pesticides (herbicides, insecticides, and fungicides) to protect crops and increase production [14,21,24].

**Fig 1:**
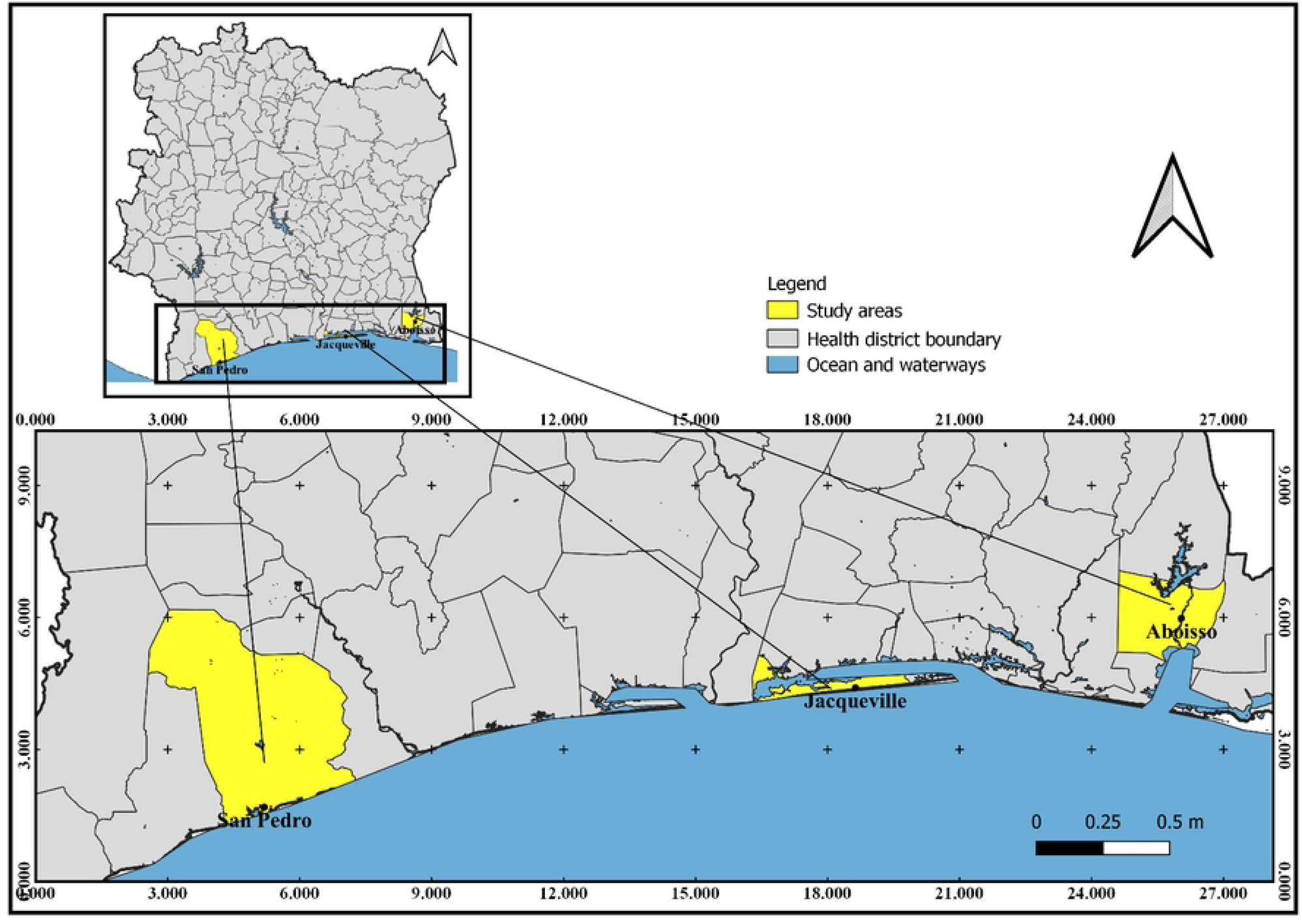
Map of Côte d’Ivoire showing the location of the littoral health district.

### Larval sampling and rearing to adults

Surveys for potential mosquito larvae habitats were conducted in diverse locations such as water puddles, vegetable cultivation sites, rice fields, and other potential larval habitats within our study sites. The larvae collected from each health district were combined, regardless of their collection site. *Anopheles gambiae s.l.* larvae were collected from breeding sites (e.g., rice fields, water wells, natural breeding sites, gardens fields (tomatoes, cabbage, carrot, lettuce)) from January 2018 to December 2020. Larvae were transferred and reared to adults under standard laboratory conditions (27 ± 2^∘^C temperature; 70 ± 10% relative humidity; 12:12 hour light: dark photoperiod). Emerged adults were provided with a cotton wool pad soaked in a 10% sugar solution. Species were identified morphologically using identification keys [25].

### WHO tube bioassay

The diagnostic dose tests of pyrethroids (alphacypermethrin, deltamethrin, and permethrin) were conducted in 2018, 2019, and 2020. The following tests were conducted exclusively in 2020. These include intensity tests (1X, 5X, and 10X), tests with the synergist (PBO), and clothianidin tests. Emerged adult females of *An. gambiae s.l.* were tested for insecticide susceptibility according to the World Health Organization (WHO) standard procedures [26]. Females of F_0_ generations aged 2–5 days were used in all the tests. Four batches of 20–25 non-blood-fed females were introduced each in four tube tests garnished with insecticide-treated filter papers, while two batches were exposed to control tubes with untreated filter papers for one hour. Mosquitoes were exposed to the WHO discriminating dosages of deltamethrin (0.05%), permethrin (0.75%) and alpha-cypermethrin (0.05%) to determine their resistance status (WHO, 2018). Moreover, synergist effects were assessed by pre-exposing mosquitoes to 5% PBO for one hour before being exposed to deltamethrin 0.05%, permethrin 0.75% and alphacypermethrin 0.05% for an additional hour. To determine the pyrethroid-resistance intensity, mosquitoes were exposed to 1X, 5X and 10X diagnostic concentrations of alpha-cypermethrin (0.05%, 0.25% and 0.5%), permethrin (0.75%, 3.75% and 7.5%) and deltamethrin (0.05%, 0.25% and 0.5%). Mosquito mortality was recorded at 24 hours after exposure.

For clothianidin, impregnated papers were prepared in *situ* following the standard operating procedure (SOP) developed by the Vector Link entomology team. The diagnostic dose was set at 2% clothianidin. Clothianidin was tested using WHO susceptibility tests, with the standard guidelines [27]. A solution of clothianidin was prepared by diluting 264 mg of the formulated product (Sumishield WG50) in 20 ml of distilled water. Four Whatman No.1 filter papers of 12 × 15 cm each were impregnated using a pipette to dispense 2 ml of solution on each filter paper, resulting in a concentration of 13.2 mg/ai clothianidin per paper. The untreated filter papers were treated using 2 ml of a solution of distilled water. Mosquitoes were tested against clothianidin-treated papers as described above [28]. The mortality was recorded daily for up to seven days in order to capture any delayed mortality effects.

### CDC bottle test

The CDC bottle tests were exclusively conducted in 2020. Chlorfenapyr susceptibility was determined using Centers of Disease Control and Prevention (CDC) bottle tests. The tests were used 250 ml glass bottles coated with 100 μg ai/bottle and 200 μg ai/bottle. A set of 15 to 20 non-blood-fed female *An. gambiae s.l.* aged 2–5 days were exposed to two discriminating concentrations of chlorfenapyr following the CDC bottle test protocol [29]. Mortality was recorded daily for up to three days.

### Molecular analyses of *An. gambiae s.l.* complex members and resistance genes

Dead and surviving mosquitoes’ exposure in WHO tube bioassays and CDC bottle tests were separately stored individually in Eppendorf tubes in silica gel and kept at −20 °C for further identification of *An*. *gambiae s.l.* complex members and *kdr* and *Ace-1* mutations.

### DNA extraction

The genomic DNA of individual mosquito was extracted according to the method described in *Collins et al.* [11]. In the Eppendorf^®^ tube (1.5 ml) individual mosquito was soaked and crushed in 200 μl of 2% Cetyl Trimethyl Ammonium Bromide (CTAB) and incubated at 65 ^∘^C for 5 min. A total of 200 μl of chloroform was added and the resulting mixture was centrifuged for 5 min at 12,000 rpm. The supernatant was pipetted into a new 1.5 ml tube. Then, 200 μl isopropanol was added, mixed by pipetting and centrifuged for 15 min at 12,000 rpm to precipitate the DNA. DNA pellet formed at the bottom of tubes. The supernatant was discarded. With 200 μl of 70% Ethanol purified the DNA of each sample and centrifuged for 5 min at 12,000 rpm. The ethanol was removed, and the pellet dried on the bench overnight. The extracted DNA was reconstituted in 30 μl DNase-free water (Sigma-Aldrich, United Kingdom) and incubated at 65 ^∘^C for 15 min and stored in the fridge at −20^∘^C.

### Complex member identification

The molecular identification of *An. gambiae* complex members was performed according to the SINE-PCR method [31], with minor modifications of reaction mixture. The PCR assay was performed using two primers, 6.1a (5’-TCGCCTTAGACCTTGCGTTA-3’) and 6.1b (5’-CGCTTCAAGAATTCGAGATAC-3’). Each reaction mixture was done in a final volume of 25 μl containing 12.5 μl One taq Quick-load 2X Master Mix, 5.5 μl free water, 0.5 μl Dimethyl sulphoxide (DMSO), 2 μl Bovine Serum Albumin (BSA), 1 μl of each primer, 0.5 μl Magnesium chloride (MgCl2) and 2 μl DNA template. The incubation took place in a thermocycler of LongGene® type (A200 Gradient Thermal cycler; LongGene Scientific Instruments Co., Ltd Hangzhou, P.R. China) according to the following program: 94 ^∘^C for 5 min, 94 ^∘^C for 25 sec, and 54 ^∘^C for 30 sec; 72 ^∘^C for 1 min repeated 35 times; and a final step at 72 ^∘^C for 10 min to terminate the reaction. After amplification, PCR products were run on either a 1.5% agarose gel in Tris/borate/EDTA (TBE) and stained with ethidium bromide solution for UV visualization. PCR products were loaded on the gel and allowed to migrate under a voltage of 140 V for one hour. The result was visualized with a UV illuminator (BioDoc-It Imaging System; Upland, CA, USA).

### Identification of target site mutation

The presence of insecticide resistance genes including *Vgsc*-L995F (previously known as *Vgsc-* L1014F (*Kdr-West*)), *Vgsc*-L995S (previously known as *Vgsc*-L1014S (*Kdr-East*)), and Ace1*-* G280S (previously known as Ace1*-*G119S)[32] was investigated using real-time PCR. The TaqMan assays as detailed by Bass [33] were used to screen for the 995F, 995S kdr and 280S mutations. The reaction was carried out in an Agilent Stratagene MX3005 qPCR thermocycler (Agilent Technologies, Santa Clara, CA, USA). For each 1 μl gene DNA, a final volume of 9 μl mix is used. The mix contains DNase-free water (3.875 μl), sensimix (5 μl) and specific primer/Probe (0.125 μl) for *kdr* 995F or 995S or *Ace1*-280S. The specific probe contains FAM and HEX fluorochromes. FAM was used to detect the mutant allele, while HEX detected the wild-type susceptible allele. PCR conditions used were 10 min at 95 ^∘^C (1 cycle), followed by 40 cycles of 10 sec at 95 ^∘^C, and 45 sec at 60 ^∘^C.

### Data analysis

The resistance status of each mosquito population was determined according to the WHO criteria with mortality after 24-hour, 72-hour and 7-day post-exposure for pyrethroids, chlorfenapyr and clothianidin, respectively [29]. Mortality was corrected using Abbott’s formula 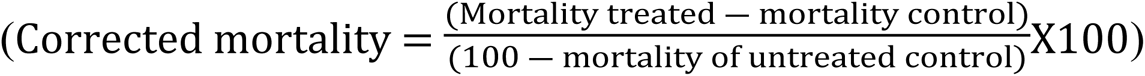, when the mortality of the control tubes was above 5% and less than 20% [34]. There was a confirmed resistance if the mortality percentage was <90%, possible resistance if the mortality rate was between 90 and 98%, and susceptibility if the mortality rate was ≥98%. The mortality recorded each year was compared between each insecticide (pyrethroid diagnostic concentration) using the ANOVA test or Kruskal-Wallis. For the resistance intensity, if corrected mortality was 98– 100% at 5X the diagnostic dose indicated low resistance intensity; if corrected mortality was less than 98% at 5X diagnostic dose implied testing the 10X diagnostic dose; if corrected mortality was 98–100% at 10X the diagnostic dose confirmed a moderate resistance intensity; if corrected mortality was less than 98% at 10X the diagnostic doses indicated high resistance intensity. For the synergist assays, the mortality of mosquitoes exposed to PBO with pyrethroid was compared with that of the insecticides without PBO pre-exposure. Comparison was made between mortality rates with and without PBO pre-exposure using the *prop .test* with software R version 4.2.1. The frequency of resistance mutations (*Vgsc*-995F, *Vgsc*-995S and *Ace1*-280S) was determined using the formula: F = [(2RR+ RS)] / [2(RR+RS+SS)] [35] with SS, homozygous susceptible genotype; RS, heterozygous genotype; RR, homozygous resistant genotype.

## Results

### Species composition of the *Anopheles gambiae* complex

A total of 672 *An. gambiae s.l.* was selected from the three sites for species identification. Overall, *An. gambiae* complex was mainly composed of *An. coluzzii* (88.2%, n=593), followed by *An. gambiae s.s.* (7.6%, n=51) and hybrids (*An. coluzzii/An. gambiae s.s.*) (4.2%, n=27). In Jacqueville and San Pedro, all mosquitoes were identified as *An. coluzzii.* The three forms were found in sympatry in Aboisso. However, *An. coluzzii* dominated in Aboisso (66.7%, n=158), followed by *An. gambiae s.s.* (21.5%, n=51) and finally hybrids (11.8%, n=28) (Fig 2).

**Fig 2:**
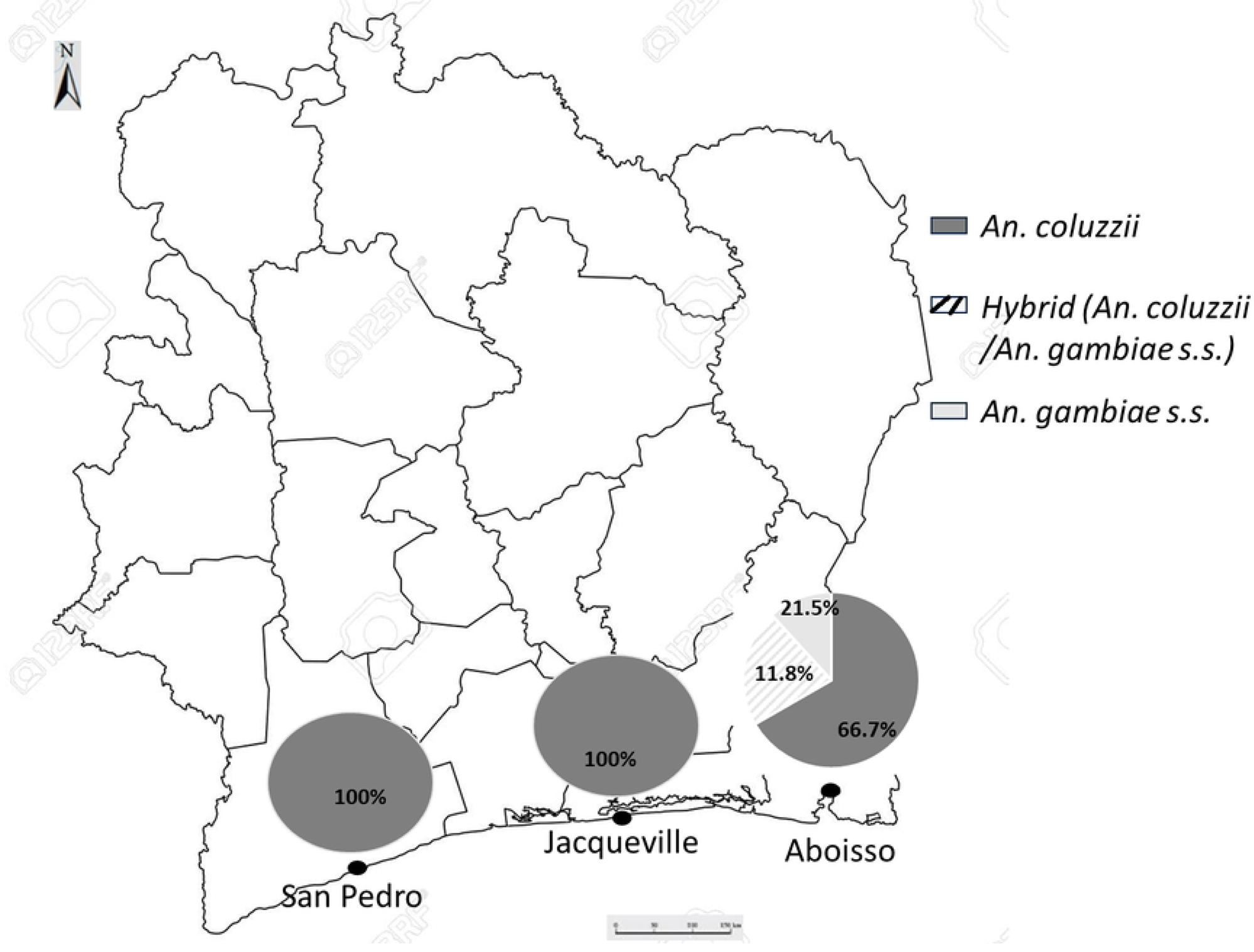
Distribution of members of *Anopheles gambiae s.l.* complex in the three study populations.

### Insecticide susceptibility in *An. gambiae s.l*

The picture (Fig 3) shows the mortality of *An. gambiae s.l.* populations from Aboisso, Jacqueville and San Pedro against alphacypermethrin, permethrin, and deltamethrin with the WHO diagnostic concentrations in 2018, 2019 and 2020.

**Fig 3:**
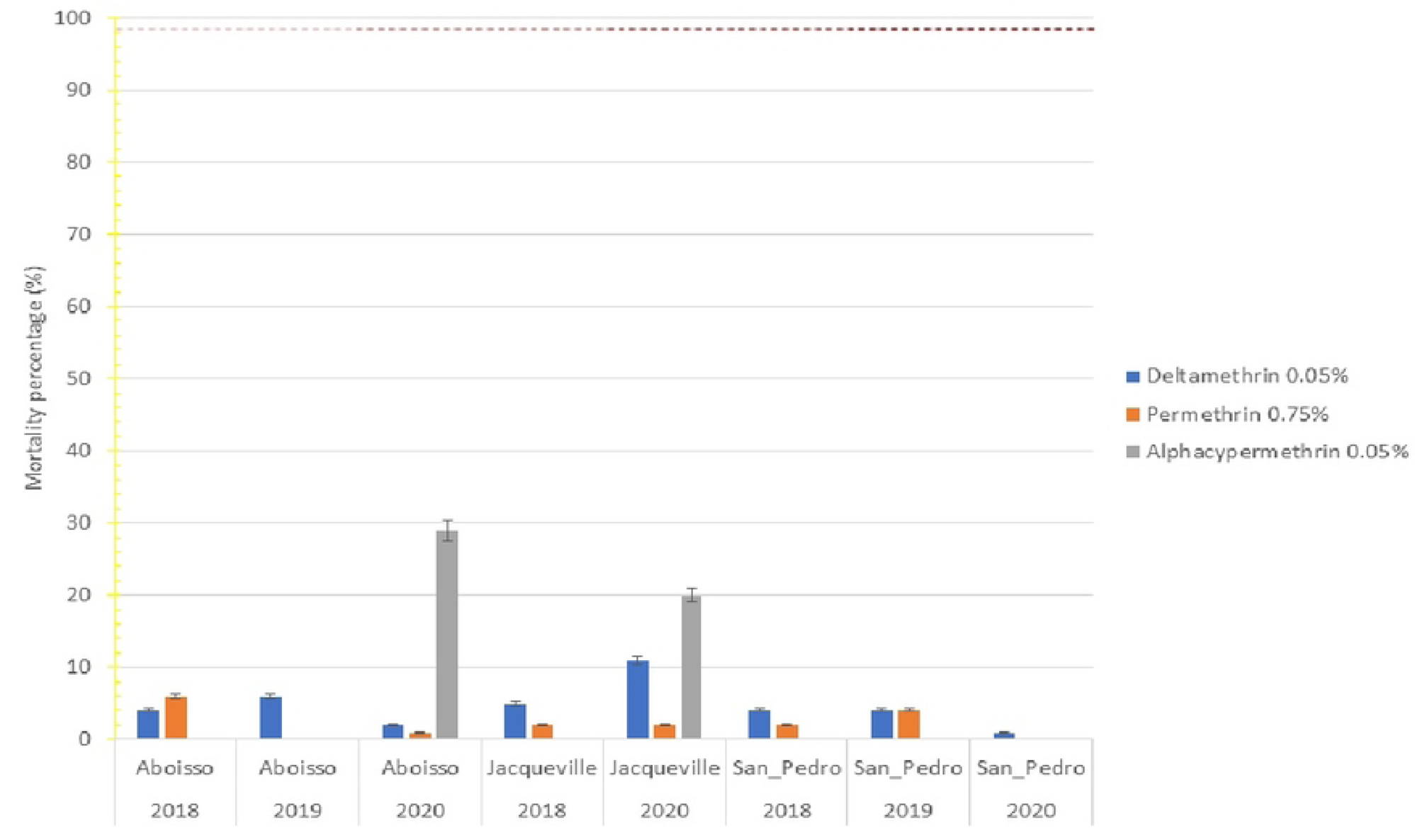
Susceptibility of *An. gambiae s.l.* to pyrethroids by site from 2018 and 2020. Error bars represent the standard errors at the Y axis and the black dashed line represents the susceptibility threshold

In 2018, in Aboisso *An. gambiae s.l.* populations, the mortality recorded was 5% for deltamethrin, 6% for permethrin and 0% for alphacypermethrin. The mortalities were not different (df = 2; p = 0.05) between insecticides. In 2019, the mortality was 6% for deltamethrin and no mortality was observed for permethrin and alphacypermethrin, the mortality rate recorded were significantly different between insecticides (df = 2; p = 0.03). The mortality recorded in 2020 were 2%, 1% and 29% respectively for deltamethrin, permethrin and alphacypermethrin. Between insecticides, the mortalities were not different (df = 2; p<0.22). In Jacqueville populations, the mortality recorded in 2018 were 5% for deltamethrin, 2% for permethrin and 0% for alphacypermethrin. The mortalities observed were not different (df = 2; p<0.07) between insecticides. In 2020, no variation was observed between each insecticide (df = 2; p = 0.12), the mortality recorded was 11% for deltamethrin, 2% for permethrin and 20% for alphacypermethrin.

In San Pedro populations, in 2018, the mortality rates were 4% for deltamethrin, 2% for permethrin and 0% for alphacypermethrin. The mortality rates recorded in 2019 were 4% for deltamethrin and permethrin and 0% for alphacypermethrin. In 2020, the mortality was 1% for permethrin and no mortality (0%) was observed for deltamethrin and alphacypermethrin. Each year, the mortality rates did not significantly differ between insecticides (Each p>0.05).

### Intensity of resistance

The results of the pyrethroid intensity test are summarized in Figure 4. These results confirmed the strong resistance of *An. gambiae s.l.* against alpha-cypermethrin, permethrin and deltamethrin. The results of resistance pyrethroid intensity tests, for all three samples from Aboisso, Jacqueville, and San Pedro, the mortalities remained below the 98% threshold when exposed to 1X and 5X concentrations of all insecticides (Fig 4). At 10X concentration, overall high intensity resistance was still observed in all three mosquito populations for the insecticides, except for moderate intensity resistance recorded with alphacypermethrin 0.5% in Aboisso populations with 100% mortality.

**Fig 4:**
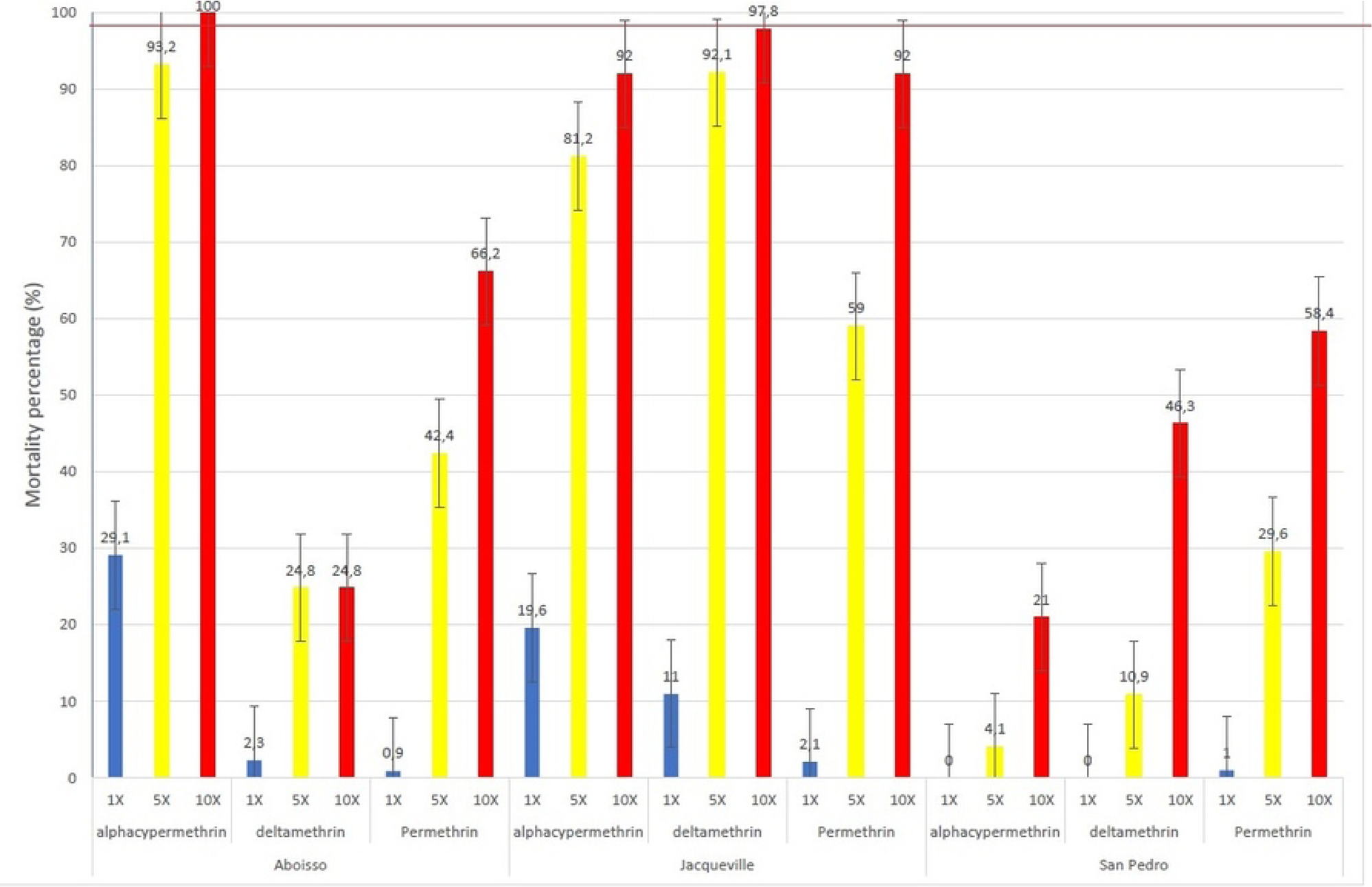
Intensity bioassays of pyrethroid against *An. gambiae s.l.* of the three study populations. Error bars represent the standard errors and the red dashed line represents the susceptibility threshold

### Effect of piperonyl butoxide (PBO)

Table 1 shows the mortality rates in the Aboisso, Jacqueville and San Pedro *An. gambiae s.l.* populations exposed to alpha-cypermethrin, permethrin and deltamethrin without and with pre-exposure to PBO. The results showed that synergist effect of PBO was strongest in all mosquito populations. Mortality increased significantly when mosquitoes were pre-exposed to PBO (df=1, p <0.0001) (Table 1). Although a significant increase was observed in mortality after pre-exposure to PBO, susceptibility was not fully restored. The mortality rates after PBO pre-exposure were still under the 98% threshold in all three study sites, except for the Jacqueville populations in which alphacypermethrin mortality after pre-exposure to PBO was 98.5% (Table 1).

**Table 1:** Bioassays of *An. gambiae s.l.* to pyrethroids without and with piperonyl butoxide of the study population.

|  |  |  |  |  |
| --- | --- | --- | --- | --- |
| Populations | Insecticide | N tested (Dead) | Mortality percentage (95% CI) | p-value |
| Aboisso | Alphacypermethrin | 82 (24) | 29.1 (19.73-40.35) | <0.0001 |
|  | PBO+Alphacypermethrin | 76 (72) | 94.7 (87.07-98.55) |  |
|  | Deltamethrin | 79 (2) | 2.3 (0.31-8.85) | <0.0001 |
|  | PBO + Deltamethrin | 84 (42) | 50.0 (38.88-61.11) |  |
|  | Permethrin | 105 (1) | 0.9 (0.02-5.19) | <0.0001 |
|  | PBO + Permethrin | 86 (28) | 32.5 (22.84-43.52) |  |
| Jacqueville | Alphacypermethrin | 89 (18) | 19.6 (12.45-30.07) | <0.0001 |
|  | PBO+Alphacypermethrin | 82 (74) | 90.1 (81.68-95.69) |  |
|  | Deltamethrin | 90 (10) | 0 (5.46-19.48) | <0.0001 |
|  | PBO + Deltamethrin | 71 (70) | 98.5 (92.40-99.96) |  |
|  | Permethrin | 79 (2) | 2.1 (0.31-8.85) | <0.0001 |
|  | PBO + Permethrin | 68 (22) | 31.7 (21.51-44.79) |  |
| San Pedro | Alphacypermethrin | 101 (0) | 0 (0-3.59) | <0.0001 |
|  | PBO+Alphacypermethrin | 106 (19) | 17.9 (11.15-26.57) |  |
|  | Deltamethrin | 94 (0) | 0 (0-3.85) | <0.0001 |
|  | PBO + Deltamethrin | 98 (28) | 28.3 (19.90-38.58) |  |
|  | Permethrin | 93 (1) | 1.1 (0.03-05.85) | <0.0001 |
|  | PBO + Permethrin | 103 (11) | 10.6 (5.45-18.31) |  |

N=represents the total number of mosquitos tested, CI=represents the confidence intervals, and p-value were calculated using the prop.test.

### Effect of clothianidin and chlorfenapyr

*Anopheles gambiae s.l.* populations from Jacqueville and San Pedro were fully susceptible to clothianidin, with mortalities of 98% (Fig 5). For the San Pedro population, 100% mortality was reached six days after exposure. In the Jacqueville population, the mortality rate was 98.8% seven days after exposure. However, for the Aboisso population, susceptibility was not fully restored after the 7-day post-exposure as the mortality rate was 55.7%.

**Fig 5:**
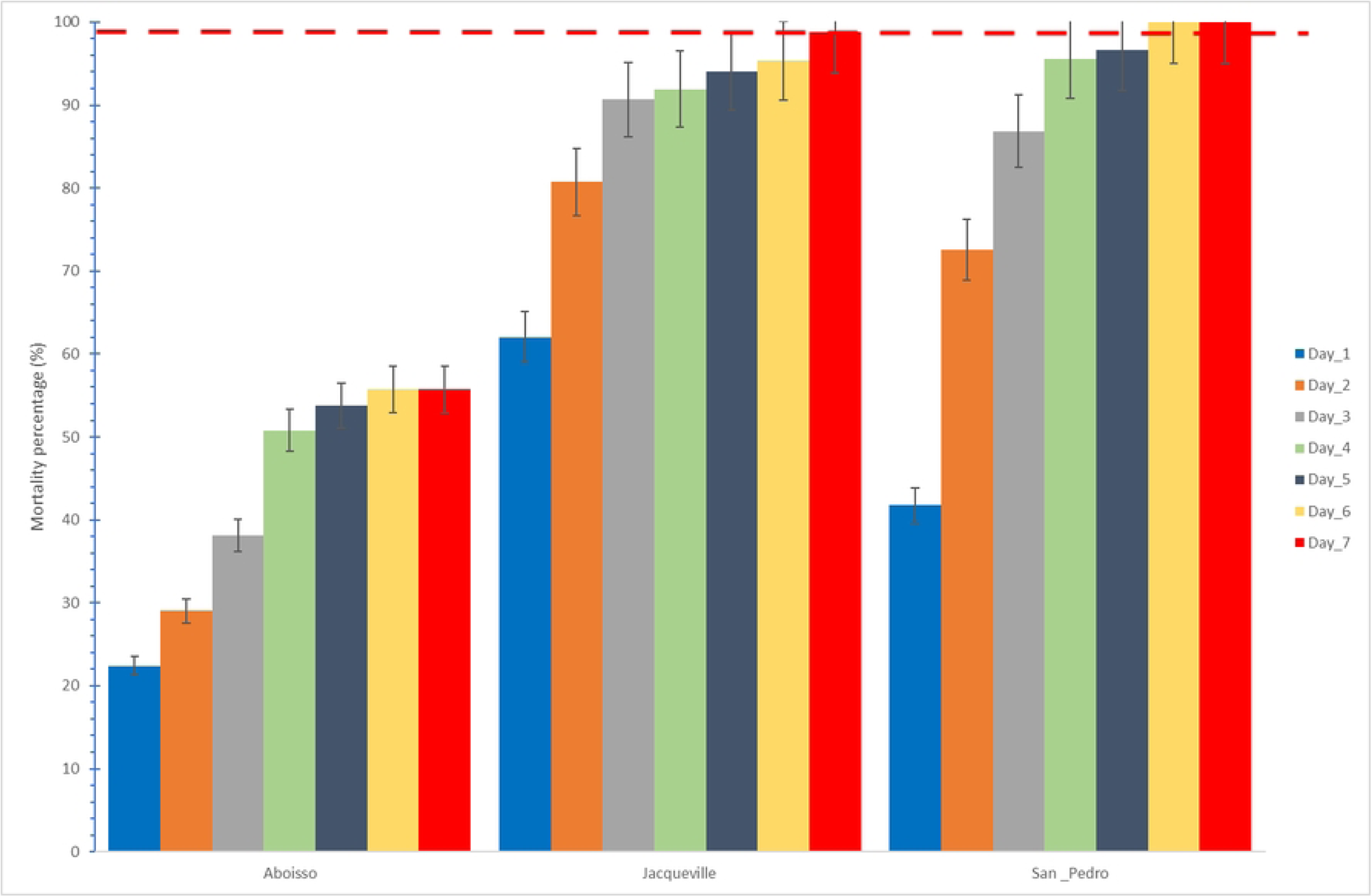
Susceptibility of *An. gambiae s.l.* to clothianidin 2% of the different populations in 2020. Error bars represent the standard errors and the red dashed line represents the susceptibility treshold.

For chlorfenapyr at 100 μg/bottle, the Aboisso population was fully susceptible, with a 72-hour mortality of 100%. However, chlorfenapyr did not fully restore susceptibility in Jacqueville and San Pedro populations with 100 μg/bottle. With 100 μg/bottle, the 72-hour mortality was of 88.7% for Jacqueville and 97.1% for San Pedro populations (Fig 6). With 200 μg/bottle, the mortality rate was 100% for the San Pedro population after 24-hours post-exposure. Except for the Jacqueville population where 100% mortality was recorded after 48-hours post-exposure (Fig 6). The population of Aboisso was not tested at 200 μg/bottle, because susceptibility was restored at 100 μg/bottle.

**Fig 6:**
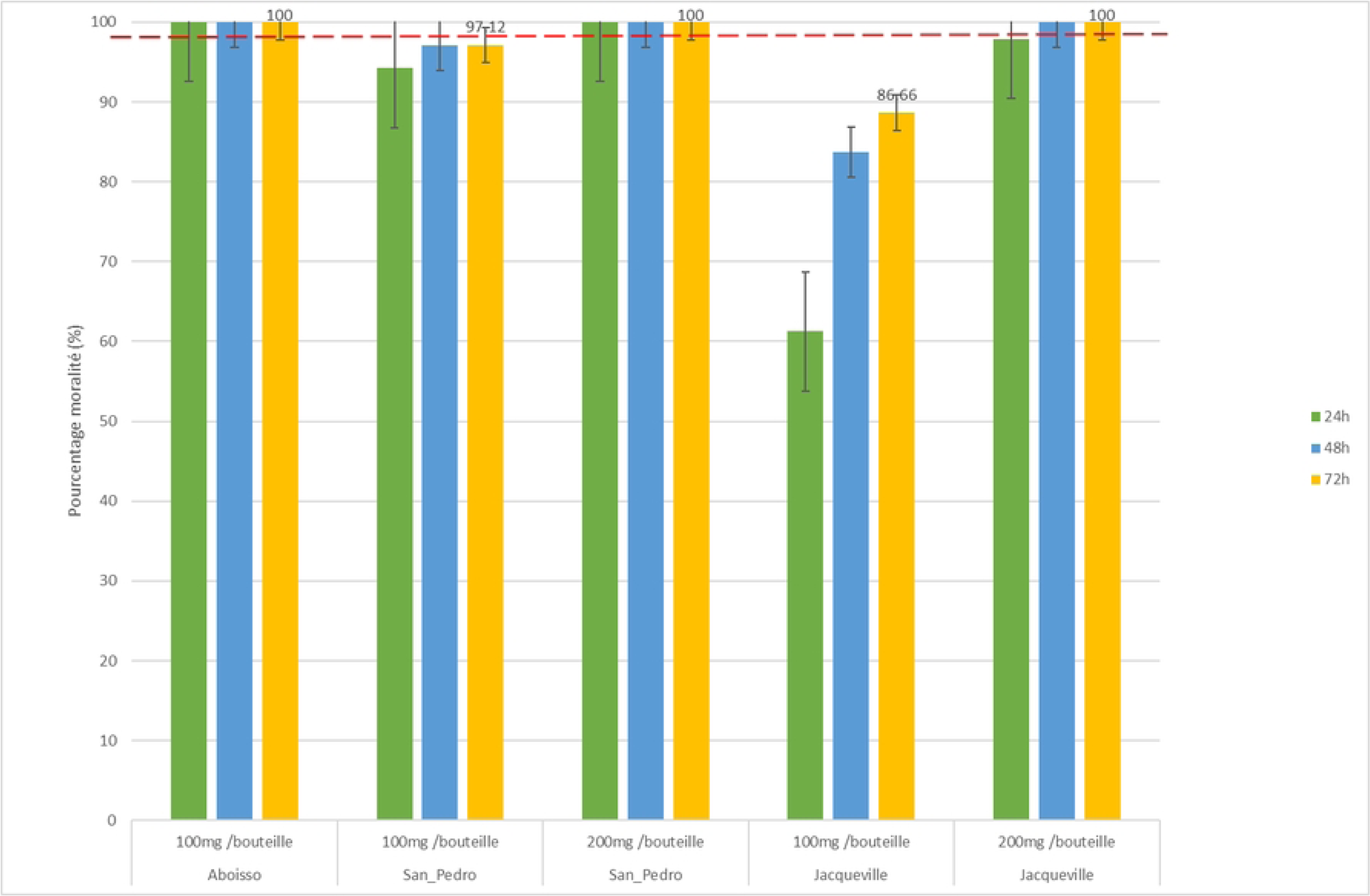
CDC bottle bioassay of chlorfenapyr against *An. gambiae s.l.* of the three populations in 2020. Error bars represent the standard errors and the red dashed line represents the susceptibility treshold.

### Resistance mutation

Table 2 shows the *kdr* (L995F and L995S) mutation allelic frequencies in the population from Aboisso, Jacqueville and San Pedro (Table 2). The *kdr* L995F mutation allelic frequencies ranged from 79.3% for *An. coluzzii*, 76.0% for hybrids, and to 72.1% for *An. gambiae s.s.* in the Aboisso population. In the population from Jacqueville and San Pedro, the *kdr* L995F (*Kdr west*) frequencies were respectively 38.1% and 71.2%. For the *Kdr* L995S (*Kdr East*), the mutation allelic frequencies were low. The mutation allelic frequencies were 2.3% for *An. gambiae s.s.* in the Aboisso population and 5.7% for the San Pedro population. For the population of Aboisso (*An. coluzzii*, and Hibryd), and San Pedro mutation was not detected. The mutation allelic frequencies of G280S were summarized in table 2. The mutation allelic frequencies of G280S were 27.4% for *An. coluzzii*, and 42.9% for *An. gambiae s.s.* in the population of Aboisso. The allelic frequencies were 31.1% and 4.2% respectively for the population of Jacqueville and San Pedro.

**Table 2:** *An. gambiae s.l.* sibling species composition and *Vgsc*-995F, *Vgsc*-995S and *Ace-1* 280S frequency in three populations.

| Table 1: *An. gambiae* s.l. sibling species composition and *vgsc-995F*, *vgsc-995S* and *Ace-1 280S* frequency in three populations |  |  |  |  |  |  |  |  |  |  |  |  |  |  |  |  |
| --- | --- | --- | --- | --- | --- | --- | --- | --- | --- | --- | --- | --- | --- | --- | --- | --- |
| Populations | *An. gambiae* s.l.<br>molecular form | *Kdr-West (L995F)* |  |  |  |  | *Kdr-East (L995S)* |  |  |  |  | *Ace-1 (G280S)* |  |  |  |  |
|  |  | Tested | FF | LF | LL | %Fr (F) | Tested | SS | LS | LL | %Fr (S) | Tested | SS | GS | GG | %Fr (S) |
| Aboisso | *An. coluzzii* | 127 | 83 | 26 | 12 | 79.3 | 127 | 0 | 0 | 127 | 0 | 31 | 5 | 7 | 19 | 27.4 |
| Aboisso | Hybrids | 25 | 16 | 6 | 3 | 76.0 | 25 | 0 | 0 | 25 | 0 | 2 | 0 | 0 | 2 | 0 |
| Aboisso | *An. gambiae* s.s. | 44 | 23 | 16 | 4 | 72.1 | 44 | 0 | 2 | 42 | 2.3 | 7 | 3 | 0 | 4 | 42.9 |
| Jacqueville | *An. coluzzii* | 145 | 86 | 45 | 14 | 38.1 | 140 | 1 | 14 | 125 | 5.7 | 82 | 4 | 43 | 35 | 31.1 |
| San Pedro | *An. coluzzii* | 99 | 57 | 27 | 15 | 71.2 | 100 | 0 | 0 | 100 | 0 | 108 | 0 | 9 | 99 | 4.2 |
| Hybrids represented *An. coluzzii*/ *An. gambiae* s.s. LL, and GG homozygous susceptible genotype (SS); LF, LS, and GS heterozygous genotype (RS); FF, and SS, homozygous resistant genotype (RR). The frequencies of resistance mutations were calculated using the formula: $F = [(2RR + RS)] / [2(RR + RS + SS)]$. | | | | | | | | | | | | | | | | |

## Discussion

This study investigated the *An. gambiae* complex members and resistance levels and mechanisms to pyrethroids (i.e., alpha-cypermethrin permethrin and deltamethrin) with and without PBO, clothianidin and chlorfenapyr in three coastal health districts of Aboisso, Jacqueville and San Pedro in southern Côte d’Ivoire. To our knowledge, this is one of the rare studies investigating both, the resistance status and complex members of malaria *Anopheles* mosquitoes in Costal Côte d’Ivoire. The current study showed that *An. gambiae s.l.* was composed of three sibling species, namely *An. coluzzii*, *An. gambiae* s.s and hybrids. Among the three sibling species *An. coluzzii* was the predominant (88.24%) species. The samples from Jacqueville and San Pedro were composed only of *An. coluzzii*. The three sibling species were represented in the Aboisso *An. gambiae s.l.* population. Among the sibling species *An. Coluzzii* (66.95%) was most abundant followed by *An. gambiae* s.s and hybrids. Moreover, *An. Gambiae s.l.* showed high phenotypic and genotypic resistance to pyrethroids, with increased mortality while pre-exposed to PBO. Overall, *An. gambiae s.l.* susceptibility was fully restored by clothianidin and chlorfenapyr throughout, except for the Aboisso population with clothianidin. *Vgsc* 995F allele frequency was very high (0.72-0.80%) in all *An. gambiae s.l.* populations. The *Vgsc* 995S was very low (0-0.06%) in all *An. gambiae s.l.* populations. For Ace-1 280S, the high frequency (0.57%) was recorded in San Pedro and the low (0) from the Aboisso hybrid population.

Species identification of *An. gambiae s.l.* showed that *An. coluzzii* constituted the major specie among *An. gambiae* complex. In San Pedro and Jacqueville, only the species *An. coluzzii* has been identified, excluding the presence of *An. gambiae s.s.* or hybrids. This selective distribution of vectors is of considerable importance for understanding malaria transmission in these regions. *Anopheles coluzzii’*s host preferences and specific breeding habitats, particularly its attraction to urban areas, may explain its dominance in these forested zones [36]. Furthermore, the geographical distribution of various *An. gambiae s.l.* species may result from local ecological variations and adaptations to specific microclimates at each site. In Aboisso, the dominance of *An. coluzzii* over *An. gambiae s.s.* and hybrids carries profound implications for malaria transmission and vector control [37]. *Anopheles coluzzii* prevalence underscores its role as the primary vector, given its affinity for human hosts and urban breeding sites, which enhance the efficiency of malaria transmission [36]. The lower prevalence of *An. gambiae s.s.* may be attributed to ecological factors or competition with *An. coluzzii* [12]. Hybrid mosquitoes, comprising 11.8% of the complex, introduce variables such as insecticide resistance and altered behavior [38,39]. Customizing vector control strategies, including the use of insecticide-treated bed nets and indoor spraying, to the dominant species, particularly *An. coluzzii*, is crucial [40]. The dominance of *An. coluzzii* species among *An. gambiae* complex is commonly reported in southern Côte d’Ivoire [7,11,12,14,17,21]. Human activities such as urban Agriculture (vegetable gardens, irrigated rice fields) could have contributed to the development of the breeding sites of *An. gambiae s.l.* [24,41]. The presence of sibling species, namely *An. coluzzii*, *An. gambiae s.s.* and hybrids, which is consistent with previous studies in the same area [12,17]. Moreover, the presence of both *An. coluzzii* and *An. gambiae s.s.*, and the hybrids in the same area found here may imply that the mating appears to be occurring between the two species.

In the bioassays conducted in this study with pyrethroid, the highest mortalities were obtained with deltamethrin. Indeed, deltamethrin and alphacypermethrin were type II pyrethroids contrary to permethrin which is type I. Type II pyrethroids are distinguished from type I pyrethroids by the presence of a cyano group at the carbon of the esterified alcohol [42]. The effect of the cyano group on insecticidal activity increases mortality [43]. In such settings, deltamethrin or alphacypermethrin used to treat nets would probably be a good choice. Several studies have shown the performance of type II pyrethroids in comparison to Type I [44]. However, the mortality varied between both insecticides. Several reasons would explain deferred mortalities induced by the same insecticide in different years. Pyrethroid resistance was detected more than 30 years ago in malaria vectors (*An. gambiae s.l.*.) in Côte d’Ivoire for the first time in vectors from the Bouaké population [45]. In this study, *An. gambiae s.l.* from Aboisso, Jacqueville and San Pedro showed very strong phenotypic resistance to pyrethroids resulting in a very low mortality at standard diagnosis doses. The result is consistent with the general trends of *An. gambiae s.l.* resistance reported by previous studies in Côte d’Ivoire [15,17] and reinforces the view that Côte d’Ivoire is a hotspot of resistance in West Africa region [13,15,17]. The phenotypic resistance to pyrethroids may imply the presence of genetic resistance involving insecticide target site modification due to the use of the same insecticides in both agriculture and public health [22–24]. The coastal zones in southern Côte d’Ivoire are devoted to large agro-industrial units (cocoa, rubber and oil palm plantations [20,22,46]) that are treated with chemicals, including pesticides and insecticides to protect the crops [14, 33–35]. Indeed, the massive and incorrect use of pesticides by untrained farmers can contribute to the development of multiple resistance of *An. gambiae s.l.* against insecticides used in public health [48].

In this study, very high allelic frequencies of the L995F *kdr* gene (>38%) were found in *An. gambiae s.l.* from the coastal health districts of Aboisso, Jacquevile and San Pedro. These high allele frequencies could reveal the widespread of L995F and kdr-West mutation within the local *An. gambiae s.l.*., as found in several regions of Côte d’Ivoire [12, 14, 15]. The low allelic frequencies of the L995S *kdr* gene (<6%) were detected in local *An. gambiae s.l.* The 995S allele was only found in *An. gambiae s.s.* from Aboisso and *An. coluzzii* from Jacqueville. Nonetheless, the presence of this allele can deliver further evidence for a pan-African propagation of the kdr resistance. This spread of the *kdr* 995F and 995S gene in *An. Gambiae s.l.* populations in littoral could compromise the effectiveness of the vector control tools that are currently in use. If the *kdr* 995F and 995S gene (against pyrethroids) spread, the efficacy of the malaria vector control tools (LLINs and IRS) was lost [49]. The high frequencies of G119S mutation in Aboisso and Jacqueville may reveal a large prevalence of this mutation in coastal areas of Côte d’Ivoire. *Ace-1* G119S allele confers resistance to organophosphates and carbamates that are commonly used in Côte d’Ivoire for crop protection [14, 35].

The present study shows that an increase in insecticide concentrations and PBO-pre-exposure significantly increased mortality in all three coastal populations of *An. gambiae s.l.* against all pyrethroids, but did not fully restore the susceptibility. Indeed, the current study showed that high intensity resistance is present in all three *An. gambiae s.l.* populations, with overall mortality being under 98% at 1X, 5X and 10X for all pyrethroid insecticides as reported by Kouassi [17] who carried out similar tests in Southern and central districts of Cote d’Ivoire. Such reduced insecticidal activity could result in lower personal protection [50], thus threatening the effectiveness of LLINs [29]. According to WHO, the vector control operational failure is likely to happen if resistance is confirmed at the 5X and especially at the 10X concentrations [29]. The various anthropogenic or natural xenobiotics present in local mosquito breeding sites and the biotic interaction could lead to the development of very high levels of resistance observed in the study [51]. The high intensity resistance recorded using only pyrethroid could be explained by high kdr mutation frequency, mixed function oxidases (MFO) activities, and an increase in esterase activity. However, moderate resistance intensity was recorded with a 10X concentration of alphacypermethrin in the Aboisso populations, and additional investigations are required for better understanding. In this study, pre-exposure to PBO showed significantly increased pyrethroid-induced mortality in all *An. gambiae s.l.* populations. However, full susceptibility was not recovered using PBO, except with the strain of Jacqueville with deltamethrin. Indeed, mortalities in PBO-pre-exposed mosquitoes tested against pyrethroids were still lower than the WHO susceptibility cut-off, as observed in other areas in Côte d’Ivoire [17,52]. Significant increase in mortality after exposure to PBO may indicate the presence of metabolic resistance (MFO, P450s, and esterases) [52], and this can compromise the effectiveness of non-PBO LLINs (e.g., Panda.Net, MagNet, PermaNet 2.0). In addition, non-restoration of full susceptibility with PBO might reduce the efficacy of PBO-treated LLINs (e.g., PermaNet 3.0) [53].

In contrast to PBO, chlorfenapyr and clothianidin restored fully the susceptibility in all populations of *An. gambiae s.l.* from Aboisso, Jacqueville and San Pedro, except for clothianidin in Aboisso population in the current study. The pyrrole chlorfenapyr and neonicotinoid clothianidin have new modes of action and are good candidates for malaria vector control in areas comprised of mutation and/or metabolic resistance to pyrethroids [54]. Indeed, after exposure to chlorfenapyr, full susceptibility was recovered in one of three districts at 100µg ai/bottle. However, when the samples were tested with higher dosages (200µg ai/bottle), the mortality increased, and susceptibility was restored in all districts. Chlorfenapyr requires activation, via cytochrome P450 monooxygenases [55]. Chlorfenapyr-treated LLINs (e.g., Interceptor G2 and PermaNet Dual) could be effective against *An. gambiae s.l.* in coastal Côte d’Ivoire [3,4,54]. Moreover, *An. gambiae s.l.* susceptibility was restored with clothianidin (100% mortality) in Jacqueville and San Pedro. As the National Malaria Strategic Plan of the National Malaria Control Programme of Côte d’Ivoire prioritizes indoor residual spraying (IRS), using clothianidin-based IRS (e.g., Fludora® Fusion) could be effective against the coastal *An. gambiae s.l.* populations in coastal Côte d’Ivoire.

## Conclusion

The present study showed that *An. gambiae s.l.* was mainly composed of *An. coluzzii*, followed by *An. gambiae* s.s and hybrids in costal Côte d’Ivoire. Local *An. gambiae s.l.* populations were highly resistant to pyrethroids and susceptible to clothianidin and chlorfenapyr. PBO pre-exposure increased the mortality, but susceptibility was not fully recovered. The coastal *An. gambiae s.l.* populations had high frequencies of target site mutation genes (kdr-West and kdr-East) and metabolic genes (Ace-1). Clothianidin and chlorfenapyr did restore full susceptibility in *An. gambiae s.l.* populations., suggesting that vector control tools treated with these new insecticides in costal Côte d’Ivoire.

## Author Contributions

**Jackson Koffi Ives Kouamé:** Conceptualization, Methodology, Investigation, Visualization, Data curation, Formal analysis, Writing – review & editing**. Ako Constant Victorien Edi :** Conceptualization, Methodology, Supervision**. Benjamin Guibehi Koudou** Conceptualization, Methodology, Supervision, Visualization, Writing – review & editing**. Ruth Marie Adjoua Kouamé:** Formal analysis, Writing – review & editing**. Yves Alain Kadio Kacou:** Investigation. **Firmain N’dri Yokoly** Writing – review & editing. Visualization**: Julien Bi Zahouli Zahouli** Methodology, Writing – review & editing**. Constant Guy N’Guessan Gbalegba :** Writing – review & editing. **David Malone :** Writing – review & editing.

## Acknowledgements

We gratefully acknowledge the contribution of technician Assamoi Jean Baptiste of the Centre Suisse de Recherches Scientifiques en Côte d’Ivoire for help and assistance during laboratory analysis. We also thank the local health district staff and a special thanks to the Programme National de Lutte contre le Paludisme Côte d’Ivoire for facilitating and participating in field data collection.

